## Supplementary figures for "Extensive re-modelling of the cell wall during the development of *Staphylococcus aureus* bacteraemia"

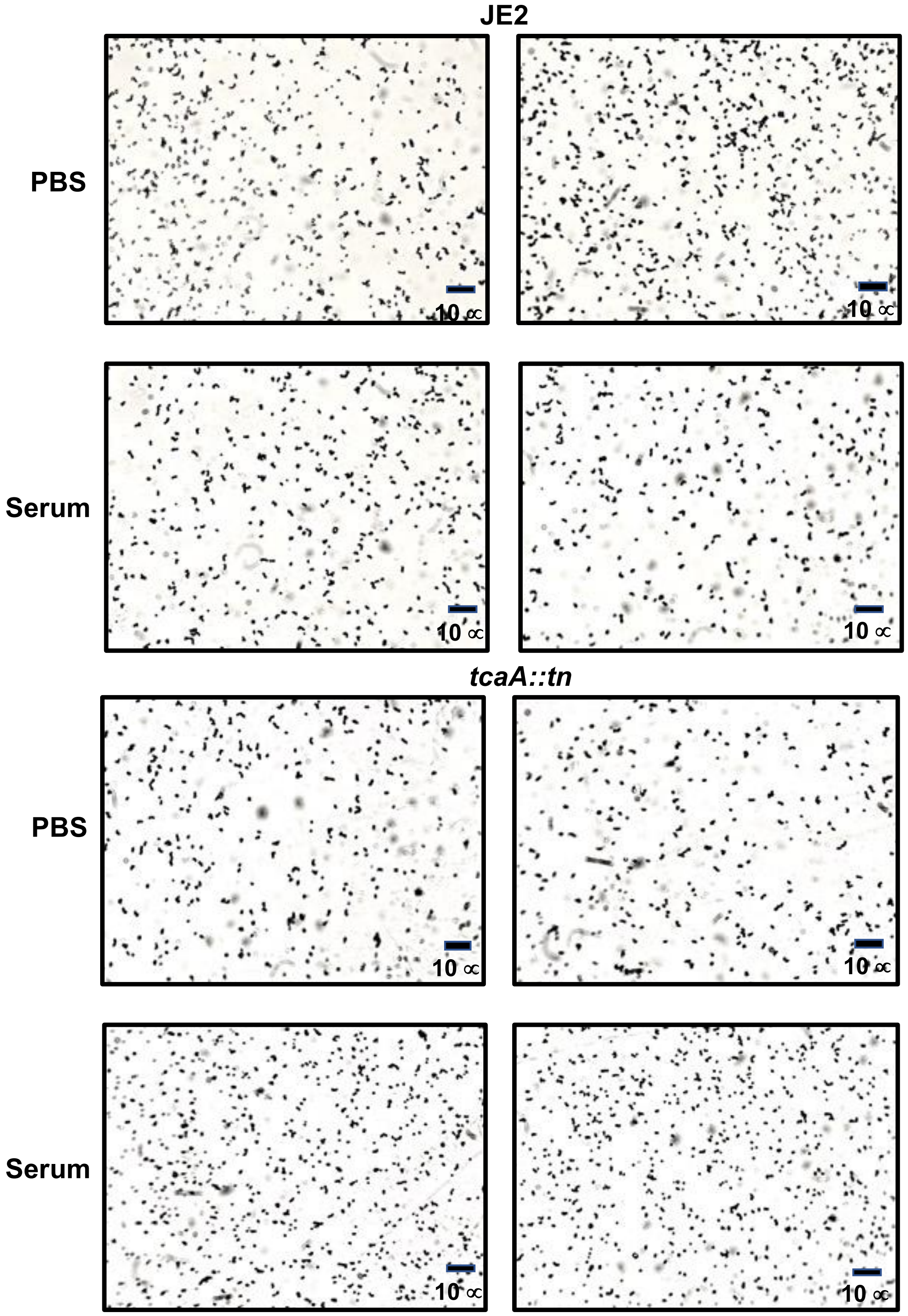


**Supp. Fig. 1:** TcaA does not contribute to the clumping of *S. aureus* when incubated with serum. A pair of light microscopy images has been provided for each of the wild type and *tcaA* mutant incubated with either PBs or human serum. No differences in clumping of the bacterial cells was observed.

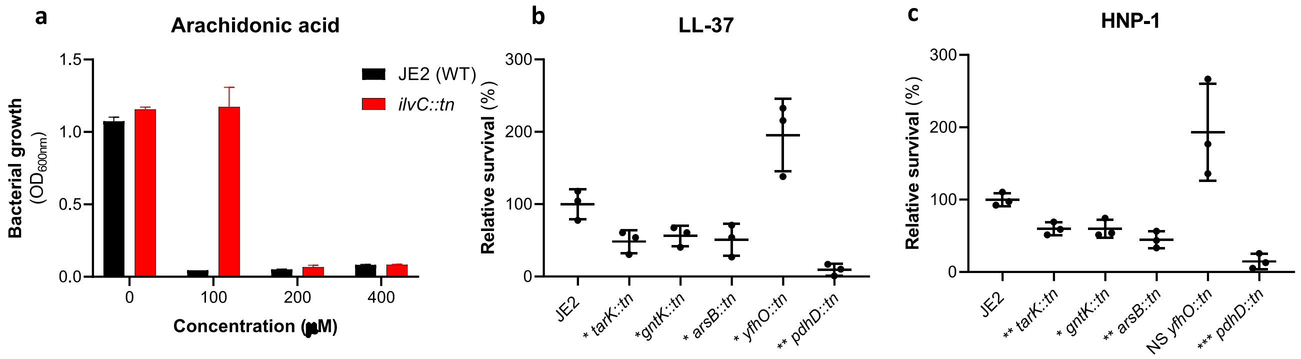


**Supp. Fig. 2:** Sensitivity of GWAS identified loci to antibacterial components of serum. **(a)** The *ilvC* mutant is more resistant to arachidonic acid. **(b & c)** the *tarK, gntK, arsB and pdhD* mutants are all more sensitive to LL-37 and HNP-1, whereas the yfhO mutant was more resistant to both.

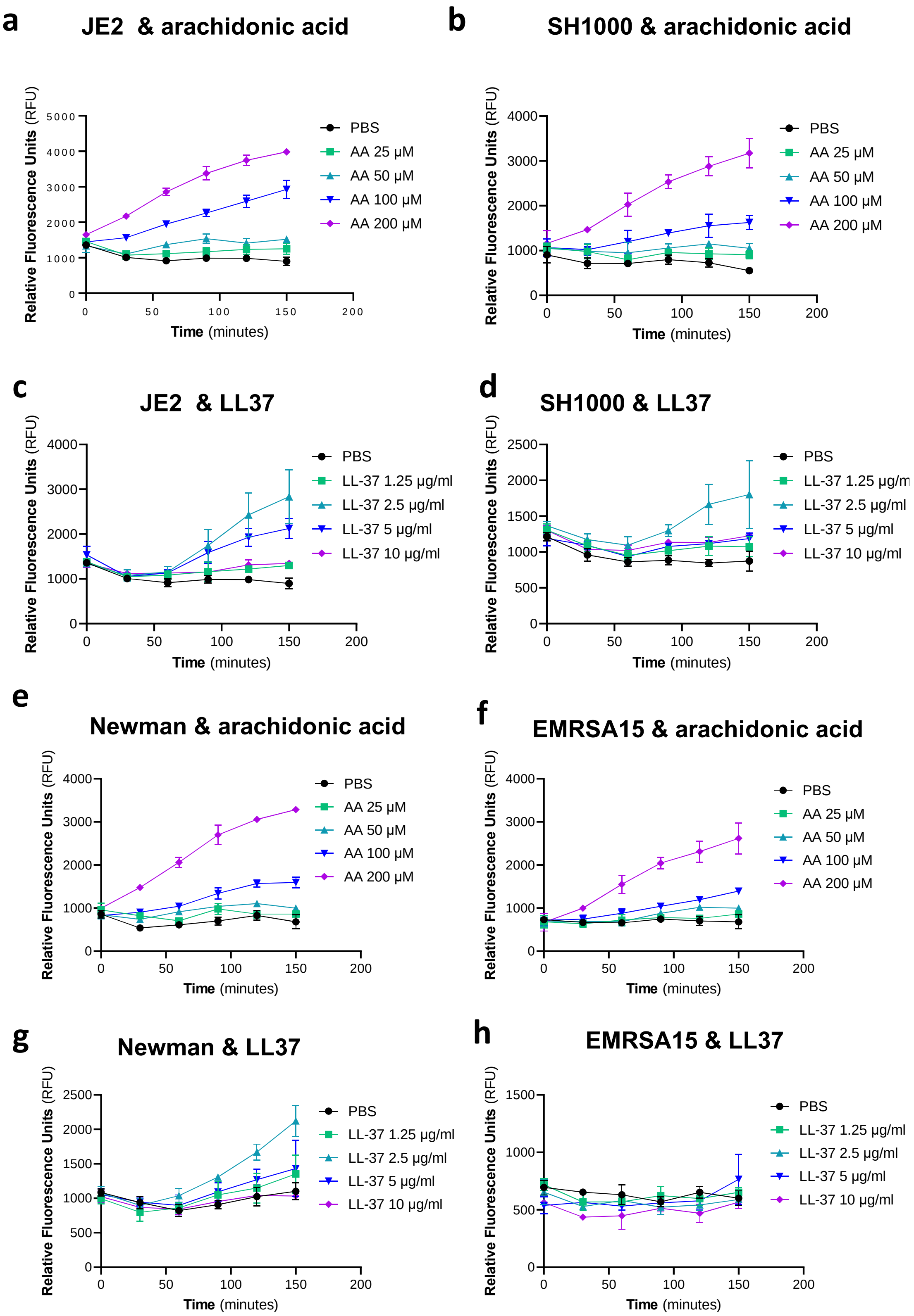


**Supp. Fig. 3:** Induction of expression of the *tcaA* gene by arachidonic acid and LL37. Using a GFP reporter system the tcaA gene eas induced in strains JE2, SH1000, Newman and EMRSA15 by arachidonic acid (a, b, e and f) and in strains JE2, SH1000 and Newman by LL37 (c, d, g). The mean of three biological replicates is presented where the error bars represent the standard error of the mean.

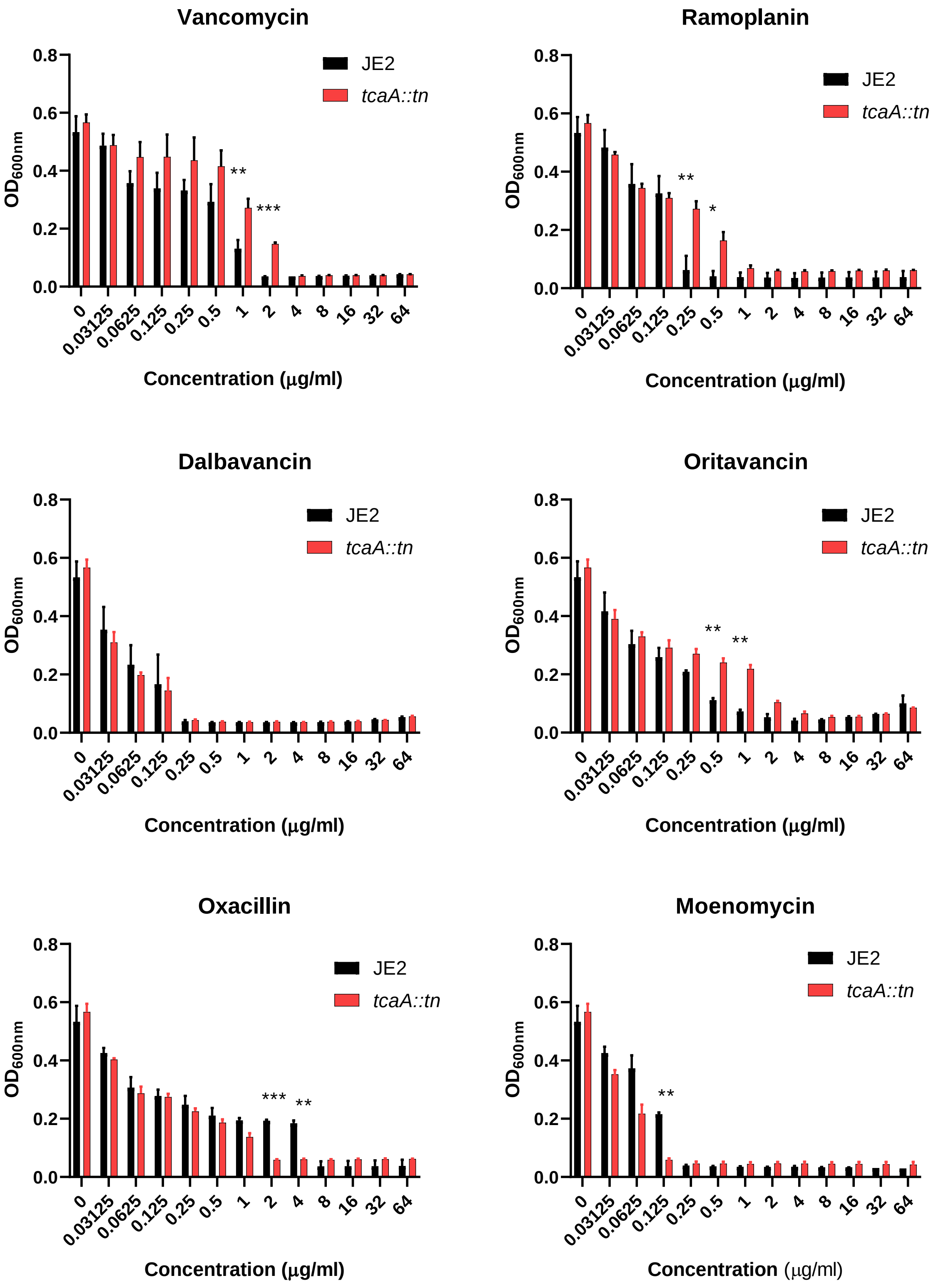


**Supp. Fig. 4:** Relative growth of the wild type JE2 and *tcaA* mutant in a range of concentrations of cell wall attacking antibiotics: vancomycin, ramoplanin, dalbavancin, oritavancin, oxacillin and moenomycin. The bars represent the mean of three biological replicates and the error bars the standard error of the mean. Significance is indicated as * p<0.05, ** p<0.01, ***p<0.001.

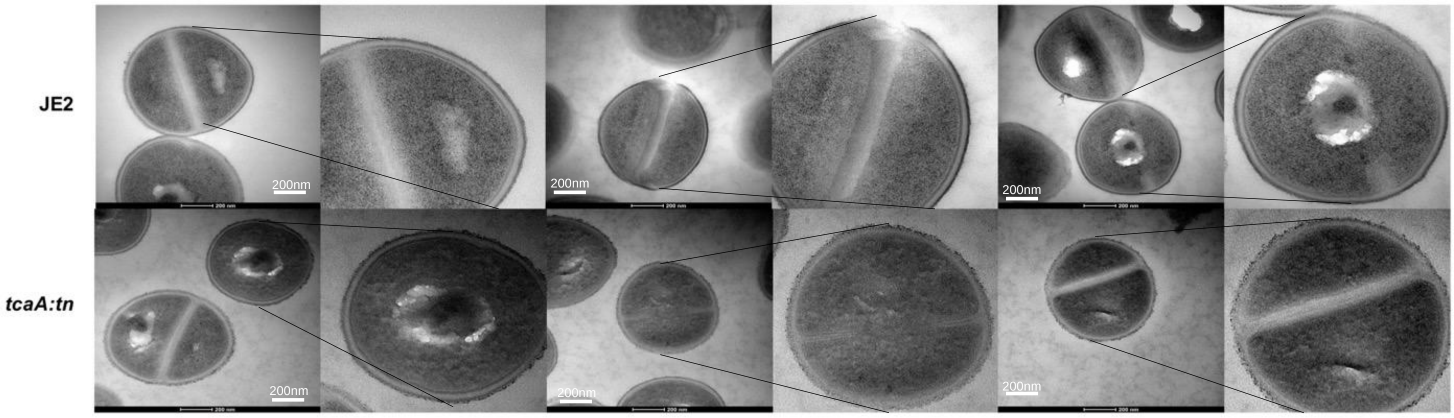


**Supp. Fig. 5:** Replicate TEM images of wild type JE2 and *tcaA* mutant cells showing the relative rough and patchy cell wall of the mutant.
