## Supplementary Table 1 for "Extensive re-modelling of the cell wall during the development of *Staphylococcus aureus* bacteraemia"

| Strain Id | CC/ST | Infection type | Outcome30days | tcaA SNP | Accession number |
|---|---|---|---|---|---|
| ASARM61 | CC22 | Bacteraemia | Alive | No | ERR084492 |
| ASARM70 | CC22 | Bacteraemia | Alive | No | ERR084501 |
| ASARM71 | CC22 | Bacteraemia | Alive | Yes | ERR084502 |
| ASARM72 | CC22 | Bacteraemia | Alive | No | ERR084503 |
| ASARM73 | CC22 | Bacteraemia | Death | No | ERR084504 |
| ASARM74 | CC22 | Bacteraemia | Death | No | ERR084505 |
| ASARM77 | CC22 | Bacteraemia | Alive | No | ERR084506 |
| ASARM76 | CC22 | Bacteraemia | Death | No | ERR084507 |
| ASARM75 | CC22 | Bacteraemia | Not known | No | ERR084508 |
| ASARM80 | CC22 | Bacteraemia | Alive | No | ERR084509 |
| ASARM79 | CC22 | Bacteraemia | Alive | No | ERR084510 |
| ASARM59 | CC22 | Bacteraemia | Alive | No | ERR084493 |
| ASARM83 | CC22 | Bacteraemia | Alive | No | ERR084513 |
| ASARM84 | CC22 | Bacteraemia | Not known | No | ERR084514 |
| ASARM86 | CC22 | Bacteraemia | Alive | No | ERR084516 |
| ASARM87 | CC22 | Bacteraemia | Alive | No | ERR084517 |
| ASARM89 | CC22 | Bacteraemia | Death | No | ERR084519 |
| ASARM62 | CC22 | Bacteraemia | Not known | No | ERR084494 |
| ASARM93 | CC22 | Bacteraemia | Alive | No | ERR084522 |
| ASARM95 | CC22 | Bacteraemia | Alive | No | ERR084523 |
| ASARM96 | CC22 | Bacteraemia | Alive | No | ERR084524 |
| ASARM97 | CC22 | Bacteraemia | Alive | No | ERR084525 |
| ASARM99 | CC22 | Bacteraemia | Death | No | ERR084527 |
| ASARM100 | CC22 | Bacteraemia | Death | No | ERR084528 |
| ASARM101 | CC22 | Bacteraemia | Death | No | ERR084529 |
| ASARM102 | CC22 | Bacteraemia | Death | No | ERR084530 |
| ASARM103 | CC22 | Bacteraemia | Death | No | ERR084531 |
| ASARM105 | CC22 | Bacteraemia | Death | No | ERR084533 |
| ASARM107 | CC22 | Bacteraemia | Alive | No | ERR084534 |
| ASARM109 | CC22 | Bacteraemia | Alive | No | ERR084535 |
| ASARM114 | CC22 | Bacteraemia | Death | No | ERR084540 |
| ASARM64 | CC22 | Bacteraemia | Alive | No | ERR109523 |
| ASARM116 | CC22 | Bacteraemia | Death | Yes | ERR084541 |
| ASARM117 | CC22 | Bacteraemia | Not known | No | ERR084542 |
| ASARM118 | CC22 | Bacteraemia | Alive | No | ERR084543 |
| ASARM119 | CC22 | Bacteraemia | Death | No | ERR084544 |
| ASARM120 | CC22 | Bacteraemia | Alive | No | ERR084545 |
| ASARM121 | CC22 | Bacteraemia | Alive | No | ERR084546 |
| ASARM122 | CC22 | Bacteraemia | Alive | No | ERR084547 |
| ASARM124 | CC22 | Bacteraemia | Alive | No | ERR084548 |
| ASARM125 | CC22 | Bacteraemia | Death | No | ERR084549 |
| ASARM126 | CC22 | Bacteraemia | Alive | No | ERR084550 |
| ASARM65 | CC22 | Bacteraemia | Alive | No | ERR084497 |
| ASARM127 | CC22 | Bacteraemia | Not known | No | ERR084551 |
| ASARM128 | CC22 | Bacteraemia | Alive | No | ERR084552 |
| ASARM129 | CC22 | Bacteraemia | Alive | No | |
| ASARM132 | CC22 | Bacteraemia | Alive | No | ERR084554 |

| | | | | | |
|---|---|---|---|---|---|
| ASARM133 | CC22 | Bacteraemia | Alive | No | ERR084556 |
| ASARM134 | CC22 | Bacteraemia | Death | No | ERR084557 |
| ASARM135 | CC22 | Bacteraemia | Alive | No | ERR084558 |
| ASARM136 | CC22 | Bacteraemia | Death | No | ERR084559 |
| ASARM137 | CC22 | Bacteraemia | Death | No | ERR084560 |
| ASARM67 | CC22 | Bacteraemia | Alive | No | ERR084498 |
| ASARM138 | CC22 | Bacteraemia | Death | No | ERR084561 |
| ASARM139 | CC22 | Bacteraemia | Alive | No | ERR084562 |
| ASARM140 | CC22 | Bacteraemia | Not known | No | ERR084563 |
| ASARM141 | CC22 | Bacteraemia | Not known | No | ERR084564 |
| ASARM142 | CC22 | Bacteraemia | Alive | No | ERR084565 |
| ASARM143 | CC22 | Bacteraemia | Alive | No | ERR084566 |
| ASARM144 | CC22 | Bacteraemia | Alive | No | ERR084567 |
| ASARM145 | CC22 | Bacteraemia | Not known | No | ERR084568 |
| ASARM68 | CC22 | Bacteraemia | Alive | No | ERR084499 |
| ASARM148 | CC22 | Bacteraemia | Not known | No | ERR084571 |
| ASARM154 | CC22 | Bacteraemia | Alive | No | ERR084575 |
| ASARM153 | CC22 | Bacteraemia | Death | No | ERR084576 |
| ASARM155 | CC22 | Bacteraemia | Alive | No | ERR084578 |
| ASARM160 | CC22 | Bacteraemia | Not known | No | ERR084579 |
| ASARM69 | CC22 | Bacteraemia | Alive | No | ERR084500 |
| ASARM162 | CC22 | Bacteraemia | Alive | No | ERR084581 |
| ASARM164 | CC22 | Bacteraemia | Not known | No | ERR084582 |
| ASARM163 | CC22 | Bacteraemia | Alive | No | ERR084583 |
| ASARM166 | CC22 | Bacteraemia | Not known | No | ERR084584 |
| ASARM165 | CC22 | Bacteraemia | Alive | No | ERR084585 |
| ASARM167 | CC22 | Bacteraemia | Alive | No | ERR084586 |
| ASARM168 | CC22 | Bacteraemia | Death | No | ERR084587 |
| ASARM169 | CC22 | Bacteraemia | Alive | No | ERR084638 |
| ASARM179 | CC22 | Bacteraemia | Death | No | ERR084648 |
| ASARM181 | CC22 | Bacteraemia | Alive | No | ERR084650 |
| ASARM183 | CC22 | Bacteraemia | Not known | No | ERR084652 |
| ASARM184 | CC22 | Bacteraemia | Alive | No | ERR084653 |
| ASARM170 | CC22 | Bacteraemia | Alive | No | ERR084639 |
| ASARM191 | CC22 | Bacteraemia | Alive | No | ERR084658 |
| ASARM193 | CC22 | Bacteraemia | Alive | No | ERR084659 |
| ASARM196 | CC22 | Bacteraemia | Alive | No | ERR084661 |
| ASARM199 | CC22 | Bacteraemia | Alive | No | ERR084664 |
| ASARM200 | CC22 | Bacteraemia | Not known | No | ERR084665 |
| ASARM201 | CC22 | Bacteraemia | Not known | No | ERR084666 |
| ASARM171 | CC22 | Bacteraemia | Alive | No | ERR084640 |
| ASARM203 | CC22 | Bacteraemia | Alive | No | ERR084667 |
| ASARM204 | CC22 | Bacteraemia | Not known | No | ERR084668 |
| ASARM205 | CC22 | Bacteraemia | Not known | No | ERR084669 |
| ASARM208 | CC22 | Bacteraemia | Alive | No | ERR084671 |
| ASARM209 | CC22 | Bacteraemia | Alive | No | ERR084672 |
| ASARM207 | CC22 | Bacteraemia | Alive | No | ERR084673 |
| ASARM211 | CC22 | Bacteraemia | Not known | No | ERR084675 |

| | | | | | |
|---|---|---|---|---|---|
| ASARM212 | CC22 | Bacteraemia | Alive | No | ERR084676 |
| ASARM172 | CC22 | Bacteraemia | Death | No | ERR084641 |
| ASARMLT1 | CC22 | Bacteraemia | Alive | No | ERR084678 |
| ASARMLT2 | CC22 | Bacteraemia | Alive | No | ERR084679 |
| ASARMLT3 | CC22 | Bacteraemia | Alive | No | ERR084680 |
| ASARM195 | CC22 | Bacteraemia | Not known | No | ERR084714 |
| ASARM176 | CC22 | Bacteraemia | Alive | No | ERR084645 |
| ASARM177 | CC22 | Bacteraemia | Death | No | ERR084646 |
| ASARM217 | CC22 | Bacteraemia | Not known | No | ERR171907 |
| ASARM220 | CC22 | Bacteraemia | Alive | No | ERR171908 |
| ASARM221 | CC22 | Bacteraemia | Alive | No | ERR171909 |
| ASARM222 | CC22 | Bacteraemia | Alive | No | ERR171910 |
| ASARM223 | CC22 | Bacteraemia | Death | No | ERR171911 |
| ASARM224 | CC22 | Bacteraemia | Alive | No | ERR171912 |
| ASARM110 | CC22 | Bacteraemia | Alive | No | ERR223125 |
| ASARM108 | CC22 | Bacteraemia | Alive | No | ERR223118 |
| ASASM42 | CC22 | Bacteraemia | Alive | No | ERR109502 |
| ASASM56 | CC22 | Bacteraemia | Death | No | ERR109515 |
| ASASM12 | CC22 | Bacteraemia | Alive | No | ERR109476 |
| ASASM61 | CC22 | Bacteraemia | Not known | No | ERR109520 |
| ASASM64 | CC22 | Bacteraemia | Not known | No | ERR084496 |
| ASASM71 | CC22 | Bacteraemia | Alive | No | ERR109528 |
| ASASM73 | CC22 | Bacteraemia | Alive | Yes | ERR109530 |
| ASASM90 | CC22 | Bacteraemia | Alive | Yes | ERR109540 |
| ASASM96 | CC22 | Bacteraemia | Not known | No | ERR109546 |
| ASASM97 | CC22 | Bacteraemia | Alive | No | ERR109547 |
| ASASM120 | CC22 | Bacteraemia | Alive | Yes | ERR109567 |
| ASASM132 | CC22 | Bacteraemia | Alive | No | ERR109578 |
| ASASM138 | CC22 | Bacteraemia | Death | No | ERR109584 |
| ASASM140 | CC22 | Bacteraemia | Alive | No | ERR109586 |
| ASASM125 | CC22 | Bacteraemia | Alive | No | ERR109572 |
| ASASM127 | CC22 | Bacteraemia | Not known | No | ERR109574 |
| ASASM181 | CC22 | Bacteraemia | Alive | Yes | ERR109623 |
| ASASM186 | CC22 | Bacteraemia | Alive | Yes | |
| ASASM190 | CC22 | Bacteraemia | Death | Yes | ERR109630 |
| ASASM246 | CC22 | Bacteraemia | Alive | No | ERR114858 |
| ASASM262 | CC22 | Bacteraemia | Alive | Yes | ERR114873 |
| ASASM390 | CC22 | Bacteraemia | Alive | Yes | ERR172029 |
| ASASM392 | CC22 | Bacteraemia | Alive | No | ERR172031 |
| ASASM398 | CC22 | Bacteraemia | Alive | Yes | ERR172036 |
| ASASM430 | CC22 | Bacteraemia | Alive | Yes | ERR172068 |
| ASASM168 | CC22 | Bacteraemia | Not known | No | ERR223120 |
| ASARM88 | CC30 | Bacteraemia | Alive | No | ERR084518 |
| ASARM90 | CC30 | Bacteraemia | Not known | No | ERR084520 |
| ASARM92 | CC30 | Bacteraemia | Alive | No | ERR084521 |
| ASARM63 | CC30 | Bacteraemia | Alive | No | ERR084495 |
| ASARM112 | CC30 | Bacteraemia | Alive | No | ERR084539 |
| ASARM147 | CC30 | Bacteraemia | Death | No | |

| | | | | | |
|---|---|---|---|---|---|
| ASARM156 | CC30 | Bacteraemia | Not known | No | ERR084577 |
| ASARM161 | CC30 | Bacteraemia | Not known | No | ERR084580 |
| ASARM180 | CC30 | Bacteraemia | Not known | No | ERR084649 |
| ASARM185 | CC30 | Bacteraemia | Not known | No | ERR084655 |
| ASARM190 | CC30 | Bacteraemia | Alive | No | ERR084657 |
| ASARM197 | CC30 | Bacteraemia | Alive | No | ERR084662 |
| ASARM210 | CC30 | Bacteraemia | Not known | No | ERR084674 |
| ASARM213 | CC30 | Bacteraemia | Alive | No | ERR084677 |
| ASASM4 | CC30 | Bacteraemia | Death | No | ERR109472 |
| ASASM22 | CC30 | Bacteraemia | Alive | No | ERR109484 |
| ASASM35 | CC30 | Bacteraemia | Alive | No | ERR109495 |
| ASASM6 | CC30 | Bacteraemia | Not known | No | ERR109474 |
| ASASM43 | CC30 | Bacteraemia | Alive | No | ERR109503 |
| ASASM46 | CC30 | Bacteraemia | Alive | No | ERR109506 |
| ASASM47 | CC30 | Bacteraemia | Alive | No | ERR109507 |
| ASASM8 | CC30 | Bacteraemia | Alive | No | ERR109475 |
| ASASM59 | CC30 | Bacteraemia | Alive | No | ERR109518 |
| ASASM14 | CC30 | Bacteraemia | Alive | No | ERR109477 |
| ASASM17 | CC30 | Bacteraemia | Alive | No | ERR109480 |
| ASASM66 | CC30 | Bacteraemia | Not known | No | ERR109525 |
| ASASM67 | CC30 | Bacteraemia | Alive | No | ERR109526 |
| ASASM78 | CC30 | Bacteraemia | Alive | No | ERR109532 |
| ASASM79 | CC30 | Bacteraemia | Alive | No | ERR109533 |
| ASASM89 | CC30 | Bacteraemia | Alive | No | ERR109539 |
| ASASM98 | CC30 | Bacteraemia | Alive | No | ERR109548 |
| ASASM99 | CC30 | Bacteraemia | Alive | No | ERR109549 |
| ASASM100 | CC30 | Bacteraemia | Not known | No | ERR109550 |
| ASASM103 | CC30 | Bacteraemia | Alive | No | ERR109553 |
| ASASM116 | CC30 | Bacteraemia | Alive | No | ERR109563 |
| ASASM131 | CC30 | Bacteraemia | Not known | Yes | ERR109577 |
| ASASM133 | CC30 | Bacteraemia | Alive | Yes | ERR109579 |
| ASASM134 | CC30 | Bacteraemia | Alive | No | ERR109580 |
| ASASM147 | CC30 | Bacteraemia | Alive | No | ERR109593 |
| ASASM149 | CC30 | Bacteraemia | Alive | No | ERR109595 |
| ASASM156 | CC30 | Bacteraemia | Alive | No | ERR109601 |
| ASASM157 | CC30 | Bacteraemia | Alive | No | ERR109602 |
| ASASM159 | CC30 | Bacteraemia | Alive | No | ERR109604 |
| ASASM166 | CC30 | Bacteraemia | Alive | No | ERR109611 |
| ASASM126 | CC30 | Bacteraemia | Death | No | ERR109573 |
| ASASM173 | CC30 | Bacteraemia | Alive | No | ERR109618 |
| ASASM174 | CC30 | Bacteraemia | Alive | No | ERR109619 |
| ASASM200 | CC30 | Bacteraemia | Alive | No | ERR109640 |
| ASASM201 | CC30 | Bacteraemia | Alive | No | ERR109641 |
| ASASM202 | CC30 | Bacteraemia | Alive | No | ERR109642 |
| ASASM212 | CC30 | Bacteraemia | Death | No | ERR109651 |
| ASASM220 | CC30 | Bacteraemia | Alive | No | ERR109658 |
| ASASM237 | CC30 | Bacteraemia | Alive | No | ERR114850 |
| ASASM250 | CC30 | Bacteraemia | Alive | No | ERR114862 |

| | | | | | |
|---|---|---|---|---|---|
| ASASM268 | CC30 | Bacteraemia | Alive | No | ERR114876 |
| ASASM272 | CC30 | Bacteraemia | Alive | No | ERR114880 |
| ASASM281 | CC30 | Bacteraemia | Death | No | ERR114887 |
| ASASM285 | CC30 | Bacteraemia | Alive | No | ERR114891 |
| ASASM288 | CC30 | Bacteraemia | Alive | No | ERR114894 |
| ASASM289 | CC30 | Bacteraemia | Alive | No | ERR114895 |
| ASASM292 | CC30 | Bacteraemia | Not known | No | ERR114898 |
| ASASM293 | CC30 | Bacteraemia | Alive | Yes | ERR114899 |
| ASASM297 | CC30 | Bacteraemia | Not known | No | ERR114902 |
| ASASM303 | CC30 | Bacteraemia | Alive | No | ERR114907 |
| ASASM307 | CC30 | Bacteraemia | Alive | Yes | ERR114910 |
| ASASM320 | CC30 | Bacteraemia | Death | No | ERR114921 |
| ASASM329 | CC30 | Bacteraemia | Alive | No | ERR114928 |
| ASASM330 | CC30 | Bacteraemia | Alive | No | ERR114929 |
| ASASM331 | CC30 | Bacteraemia | Alive | No | ERR114930 |
| ASASM340 | CC30 | Bacteraemia | Alive | No | ERR109667 |
| ASASM345 | CC30 | Bacteraemia | Alive | No | ERR109672 |
| ASASM351 | CC30 | Bacteraemia | Death | No | ERR109678 |
| ASASM358 | CC30 | Bacteraemia | Alive | No | ERR109685 |
| ASASM369 | CC30 | Bacteraemia | Not known | No | ERR109696 |
| ASASM391 | CC30 | Bacteraemia | Alive | No | ERR172030 |
| ASASM395 | CC30 | Bacteraemia | Not known | No | ERR172033 |
| ASASM406 | CC30 | Bacteraemia | Alive | No | ERR172044 |
| ASASM407 | CC30 | Bacteraemia | Alive | No | ERR172045 |
| ASASM408 | CC30 | Bacteraemia | Death | No | ERR172046 |
| ASASM436 | CC30 | Bacteraemia | Death | No | ERR172074 |
| ASASM443 | CC30 | Bacteraemia | Alive | No | ERR172081 |
| ASASM445 | CC30 | Bacteraemia | Alive | No | ERR172083 |
| ASASM446 | CC30 | Bacteraemia | Death | No | ERR172084 |
| ASASM451 | CC30 | Bacteraemia | Alive | No | ERR172089 |
| ASASM457 | CC30 | Bacteraemia | Death | No | ERR172093 |
| ASASM458 | CC30 | Bacteraemia | Alive | No | ERR172094 |
| ASASM373 | CC30 | Bacteraemia | Alive | Yes | ERR223174 |
| ASASM377 | CC30 | Bacteraemia | Not known | Yes | ERR223177 |
| ASASM378 | CC30 | Bacteraemia | Alive | No | ERR223178 |
| ASASM379 | CC30 | Bacteraemia | Alive | No | ERR223179 |
| ASASM385 | CC30 | Bacteraemia | Alive | No | ERR355917 |
| EOE003 | CC30 | Bacteraemia | Not known | No | ERR156428 |
| EOE023 | CC30 | Bacteraemia | Not known | No | ERR156447 |
| EOE030 | CC30 | Bacteraemia | Not known | No | ERR156452 |
| EOE035 | CC30 | Bacteraemia | Not known | No | ERR156457 |
| EOE041 | CC30 | Bacteraemia | Not known | No | ERR156462 |
| EOE042 | CC30 | Bacteraemia | Not known | No | ERR156463 |
| EOE045 | CC30 | Bacteraemia | Not known | No | ERR156465 |
| EOE052 | CC30 | Bacteraemia | Not known | No | ERR156470 |
| EOE054 | CC30 | Bacteraemia | Not known | No | ERR156472 |
| EOE057 | CC30 | Bacteraemia | Not known | No | ERR156473 |
| EOE061 | CC30 | Bacteraemia | Not known | No | ERR156477 |

| | | | | | |
|---|---|---|---|---|---|
| EOE065 | CC30 | Bacteraemia | Not known | No | ERR156480 |
| EOE072 | CC30 | Bacteraemia | Not known | No | ERR156486 |
| EOE073 | CC30 | Bacteraemia | Not known | No | ERR156487 |
| EOE078 | CC30 | Bacteraemia | Not known | No | ERR156493 |
| EOE083 | CC30 | Bacteraemia | Not known | No | ERR156498 |
| EOE084 | CC30 | Bacteraemia | Not known | No | ERR156499 |
| EOE086 | CC30 | Bacteraemia | Not known | No | ERR156500 |
| EOE088 | CC30 | Bacteraemia | Not known | No | ERR156502 |
| EOE089 | CC30 | Bacteraemia | Not known | No | ERR156503 |
| EOE090 | CC30 | Bacteraemia | Not known | No | ERR156504 |
| EOE091 | CC30 | Bacteraemia | Not known | No | ERR156505 |
| EOE094 | CC30 | Bacteraemia | Not known | No | ERR156507 |
| EOE096 | CC30 | Bacteraemia | Not known | No | ERR156509 |
| EOE097 | CC30 | Bacteraemia | Not known | No | ERR156510 |
| EOE098 | CC30 | Bacteraemia | Not known | No | ERR156511 |
| EOE099 | CC30 | Bacteraemia | Not known | No | ERR156512 |
| EOE100 | CC30 | Bacteraemia | Not known | No | ERR156513 |
| EOE101 | CC30 | Bacteraemia | Not known | No | ERR156514 |
| EOE102 | CC30 | Bacteraemia | Not known | No | ERR156515 |
| EOE103 | CC30 | Bacteraemia | Not known | No | ERR156516 |
| EOE104 | CC30 | Bacteraemia | Not known | No | ERR156517 |
| EOE105 | CC30 | Bacteraemia | Not known | No | ERR156518 |
| EOE106 | CC30 | Bacteraemia | Not known | No | ERR156519 |
| EOE122 | CC30 | Bacteraemia | Not known | No | ERR158987 |
| EOE125 | CC30 | Bacteraemia | Not known | Yes | ERR158990 |
| EOE129 | CC30 | Bacteraemia | Not known | No | ERR158994 |
| EOE130 | CC30 | Bacteraemia | Not known | Yes | ERR158995 |
| EOE137 | CC30 | Bacteraemia | Not known | No | ERR159002 |
| EOE140 | CC30 | Bacteraemia | Not known | No | ERR159005 |
| EOE154 | CC30 | Bacteraemia | Not known | No | ERR159018 |
| EOE155 | CC30 | Bacteraemia | Not known | No | ERR159019 |
| EOE158 | CC30 | Bacteraemia | Not known | No | ERR159022 |
| EOE118 | CC30 | Bacteraemia | Not known | No | ERR158982 |
| EOE198 | CC30 | Bacteraemia | Not known | No | ERR159048 |
| EOE205 | CC30 | Bacteraemia | Not known | No | ERR159053 |
| EOE220 | CC30 | Bacteraemia | Not known | No | ERR159065 |
| EOE120 | CC30 | Bacteraemia | Not known | No | ERR158985 |
| EOE225 | CC30 | Bacteraemia | Not known | No | ERR159070 |
| EOE233 | CC30 | Bacteraemia | Not known | No | ERR177157 |
| EOE234 | CC30 | Bacteraemia | Not known | No | ERR177158 |
| EOE237 | CC30 | Bacteraemia | Not known | No | ERR177162 |
| EOE268 | CC30 | Bacteraemia | Not known | No | ERR177193 |
| EOE269 | CC30 | Bacteraemia | Not known | No | ERR177194 |
| EOE274 | CC30 | Bacteraemia | Not known | No | ERR177199 |
| EOE275 | CC30 | Bacteraemia | Not known | No | ERR177200 |
| EOE276 | CC30 | Bacteraemia | Not known | No | ERR177201 |
| EOE029 | CC30 | Bacteraemia | Not known | No | ERR177204 |
| EOE161 | CC30 | Bacteraemia | Not known | No | ERR177208 |

| | | | | | |
|---|---|---|---|---|---|
| EOE162 | CC30 | Bacteraemia | Not known | No | ERR177209 |
| EOE163 | CC30 | Bacteraemia | Not known | No | ERR177211 |
| EOE165 | CC30 | Bacteraemia | Not known | No | ERR177212 |
| EOE166 | CC30 | Bacteraemia | Not known | No | ERR177213 |
| EOE167 | CC30 | Bacteraemia | Not known | No | ERR177214 |
| EOE169 | CC30 | Bacteraemia | Not known | No | ERR177215 |
| EOE170 | CC30 | Bacteraemia | Not known | No | |
| EOE171 | CC30 | Bacteraemia | Not known | No | ERR177217 |
| EOE173 | CC30 | Bacteraemia | Not known | No | ERR177218 |
| EOE174 | CC30 | Bacteraemia | Not known | No | ERR177219 |
| EOE175 | CC30 | Bacteraemia | Not known | No | ERR177220 |
| EOE176 | CC30 | Bacteraemia | Not known | No | ERR177221 |
| EOE208 | CC30 | Bacteraemia | Not known | No | ERR177225 |
| EOE126 | CC30 | Bacteraemia | Not known | No | ERR355922 |
| EOE229 | CC30 | Bacteraemia | Not known | No | ERR355923 |
| Sa_TPS3105 | ST93 | Bacteraemia | Not known | No | |
| Sa_TPS3148 | ST93 | Bacteraemia | Not known | No | |
| Sa_TPS3161 | ST93 | SSTI | Not known | No | |
| Sa_TPS3151 | ST93 | SSTI | Not known | No | |
| Sa_TPS3165 | ST93 | Carriage | Not known | No | |
| Sa_TPS3150 | ST93 | Sputum | Not known | No | |
| Sa_TPS3171 | ST93 | Carriage | Not known | No | |
| Sa_TPS3155 | ST93 | Carriage | Not known | No | |
| Sa_TPS3118 | ST93 | not recorded | Not known | No | |
| Sa_TPS3167 | ST93 | ndotrachael tub | Not known | No | |
| Sa_TPS3162 | ST93 | Carriage | Not known | No | |
| Sa_TPS3169 | ST93 | SSTI | Not known | No | |
| Sa_TPS3106 | ST93 | Carriage | Not known | No | |
| Sa_TPS3145 | ST93 | SSTI | Not known | No | |
| Sa_TPS3158 | ST93 | SSTI | Not known | No | |
| Sa_TPS3183 | ST93 | not recorded | Not known | No | |
| Sa_TPS3134 | ST93 | SSTI | Not known | No | |
| Sa_TPS3137 | ST93 | SSTI | Not known | No | |
| Sa_TPS3181 | ST93 | SSTI | Not known | No | |
| Sa_TPS3026 | ST93 | Bacteraemia | Not known | No | |
| Sa_TPS3142 | ST93 | Bacteraemia | Not known | No | |
| Sa_TPS3164 | ST93 | Carriage | Not known | No | |
| Sa_TPS3153 | ST93 | Carriage | Not known | No | |
| Sa_TPS3104 | ST93 | Carriage | Not known | No | |
| Sa_TPS3139 | ST93 | SSTI | Not known | No | |
| Sa_TPS3149 | ST93 | Carriage | Not known | No | |
| Sa_TPS3138 | ST93 | SSTI | Not known | No | |
| Sa_TPS3160 | ST93 | SSTI | Not known | No | |
| Sa_TPS3185 | ST93 | Carriage | Not known | No | |
| Sa_TPS3176 | ST93 | SSTI | Not known | No | |
| Sa_TPS3144 | ST93 | SSTI | Not known | No | |
| Sa_TPS3174 | ST93 | SSTI | Not known | No | |
| Sa_TPS3146 | ST93 | SSTI | Not known | No | |

| | | | | |
|---|---|---|---|---|
| Sa_TPS3133 | ST93 | Carriage | Not known | No |
| Sa_TPS3132 | ST93 | not recorded | Not known | No |
| Sa_TPS3152 | ST93 | pneumoniae | Not known | No |
| Sa_TPS3135 | ST93 | Carriage | Not known | No |
| Sa_TPS3173 | ST93 | Carriage | Not known | No |
| Sa_TPS3140 | ST93 | SSTI | Not known | No |
| Sa_TPS3136 | ST93 | Bacteraemia | Not known | No |
| Sa_TPS3188 | ST93 | not recorded | Not known | No |
| Sa_TPS3166 | ST93 | Carriage | Not known | No |
| Sa_TPS3178 | ST93 | Pleural fluid | Not known | No |
| Sa_TPS3157 | ST93 | SSTI | Not known | No |
| Sa_TPS3168 | ST93 | SSTI | Not known | No |
| Sa_TPS3189 | ST93 | not recorded | Not known | No |
| Sa_TPS3154 | ST93 | SSTI | Not known | No |
| Sa_TPS3163 | ST93 | SSTI | Not known | No |
| Sa_TPS3184 | ST93 | Carriage | Not known | No |
| Sa_TPS3182 | ST93 | not recorded | Not known | No |
| Sa_TPS3147 | ST93 | SSTI | Not known | No |
| Sa_TPS3177 | ST93 | Bacteraemia | Not known | Yes |
| Sa_TPS3159 | ST93 | SSTI | Not known | No |
| Sa_TPS3156 | ST93 | Carriage | Not known | No |
| Sa_TPS3180 | ST93 | not recorded | Not known | No |
| Sa_TPS3186 | ST93 | Carriage | Not known | No |
| Sa_TPS3179 | ST93 | Carriage | Not known | No |
| Sa_TPS3187 | ST93 | Carriage | Not known | No |
| MR005 | ST8 | Bacteraemia | Not known | No |
| MR007 | ST8 | Bacteraemia | Not known | No |
| MR018 | ST8 | Bacteraemia | Not known | No |
| MR019 | ST8 | Bacteraemia | Not known | No |
| MR021 | ST8 | Bacteraemia | Not known | No |
| MR022 | ST8 | Bacteraemia | Not known | No |
| MR023 | ST8 | Bacteraemia | Not known | No |
| MR025 | ST8 | Bacteraemia | Not known | No |
| MR026 | ST8 | Bacteraemia | Not known | No |
| MR027 | ST8 | Bacteraemia | Not known | No |
| MR029 | ST8 | Bacteraemia | Not known | No |
| MR030 | ST8 | Bacteraemia | Not known | No |
| MR031 | ST8 | Bacteraemia | Not known | No |
| MR035 | ST8 | Bacteraemia | Not known | No |
| MR036 | ST8 | Bacteraemia | Not known | No |
| MR039 | ST8 | Bacteraemia | Not known | No |
| MR047 | ST8 | Bacteraemia | Not known | No |
| MR051 | ST8 | Bacteraemia | Not known | No |
| MR060 | ST8 | Bacteraemia | Not known | No |
| MR063 | ST8 | Bacteraemia | Not known | No |
| MR064 | ST8 | Bacteraemia | Not known | No |
| MR065 | ST8 | Bacteraemia | Not known | No |
| MR072 | ST8 | Bacteraemia | Not known | No |

| | | | | |
|---|---|---|---|---|
| MR073 | ST8 | Bacteraemia | Not known | No |
| MR074 | ST8 | Bacteraemia | Not known | No |
| MR077 | ST8 | Bacteraemia | Not known | No |
| MR078 | ST8 | Bacteraemia | Not known | No |
| MR081 | ST8 | Bacteraemia | Not known | No |
| MR083 | ST8 | Bacteraemia | Not known | No |
| MR084 | ST8 | Bacteraemia | Not known | No |
| MR087 | ST8 | Bacteraemia | Not known | No |
| MR090 | ST8 | Bacteraemia | Not known | No |
| MR091 | ST8 | Bacteraemia | Not known | No |
| MR096 | ST8 | Bacteraemia | Not known | No |
| MR107 | ST8 | Bacteraemia | Not known | No |
| MR110 | ST8 | Bacteraemia | Not known | No |
| USFL008 | ST8 | Carriage | Not known | No |
| USFL009 | ST8 | Carriage | Not known | No |
| USFL012 | ST8 | Carriage | Not known | Yes |
| USFL028 | ST8 | Carriage | Not known | No |
| USFL042 | ST8 | Carriage | Not known | No |
| USFL061 | ST8 | Carriage | Not known | No |
| USFL063 | ST8 | Carriage | Not known | No |
| USFL074 | ST8 | Carriage | Not known | No |
| USFL077 | ST8 | Carriage | Not known | No |
| USFL082 | ST8 | Carriage | Not known | No |
| USFL093 | ST8 | Carriage | Not known | No |
| USFL119 | ST8 | Carriage | Not known | No |
| USFL130 | ST8 | Carriage | Not known | No |
| USFL141 | ST8 | Carriage | Not known | No |
| USFL153 | ST8 | Carriage | Not known | No |
| USFL156 | ST8 | Carriage | Not known | No |
| USFL166 | ST8 | Carriage | Not known | No |
| USFL167 | ST8 | Carriage | Not known | No |
| USFL169 | ST8 | Carriage | Not known | No |
| USFL182 | ST8 | Carriage | Not known | No |
| USFL200 | ST8 | Carriage | Not known | No |
| USFL211 | ST8 | Carriage | Not known | No |
| USFL213 | ST8 | Carriage | Not known | No |
| USFL224 | ST8 | Carriage | Not known | No |
| USFL225 | ST8 | Carriage | Not known | No |
| USFL230 | ST8 | Carriage | Not known | No |
| USFL231 | ST8 | Carriage | Not known | No |
| USFL243 | ST8 | Carriage | Not known | No |
| USFL248 | ST8 | Carriage | Not known | No |
| USFL259 | ST8 | Carriage | Not known | No |
| USFL263 | ST8 | Carriage | Not known | No |
| USFL267 | ST8 | Carriage | Not known | No |
| USFL269 | ST8 | Carriage | Not known | No |
| USFL271 | ST8 | Carriage | Not known | No |
| USFL272 | ST8 | Carriage | Not known | No |

| | | | | |
|---|---|---|---|---|
| USFL302 | ST8 | Carriage | Not known | No |
| USFL303 | ST8 | Carriage | Not known | No |
| USFL304 | ST8 | Carriage | Not known | No |
| USFL016 | ST8 | SSTI | Not known | No |
| USFL018 | ST8 | SSTI | Not known | No |
| USFL020 | ST8 | SSTI | Not known | No |
| USFL021 | ST8 | SSTI | Not known | Yes |
| USFL034 | ST8 | SSTI | Not known | No |
| USFL035 | ST8 | SSTI | Not known | No |
| USFL036 | ST8 | SSTI | Not known | No |
| USFL039 | ST8 | SSTI | Not known | No |
| USFL056 | ST8 | SSTI | Not known | No |
| USFL057 | ST8 | SSTI | Not known | No |
| USFL059 | ST8 | SSTI | Not known | No |
| USFL069 | ST8 | SSTI | Not known | No |
| USFL095 | ST8 | SSTI | Not known | No |
| USFL097 | ST8 | SSTI | Not known | No |
| USFL103 | ST8 | SSTI | Not known | No |
| USFL110 | ST8 | SSTI | Not known | No |
| USFL111 | ST8 | SSTI | Not known | No |
| USFL113 | ST8 | SSTI | Not known | No |
| USFL136 | ST8 | SSTI | Not known | No |
| USFL137 | ST8 | SSTI | Not known | No |
| USFL138 | ST8 | SSTI | Not known | No |
| USFL139 | ST8 | SSTI | Not known | No |
| USFL149 | ST8 | SSTI | Not known | No |
| USFL152 | ST8 | SSTI | Not known | No |
| USFL158 | ST8 | SSTI | Not known | No |
| USFL159 | ST8 | SSTI | Not known | No |
| USFL160 | ST8 | SSTI | Not known | No |
| USFL162 | ST8 | SSTI | Not known | No |
| USFL164 | ST8 | SSTI | Not known | No |
| USFL165 | ST8 | SSTI | Not known | No |
| USFL173 | ST8 | SSTI | Not known | No |
| USFL174 | ST8 | SSTI | Not known | No |
| USFL194 | ST8 | SSTI | Not known | No |
| USFL198 | ST8 | SSTI | Not known | No |
| USFL218 | ST8 | SSTI | Not known | No |
| USFL219 | ST8 | SSTI | Not known | No |
| USFL220 | ST8 | SSTI | Not known | No |
| USFL221 | ST8 | SSTI | Not known | No |
| USFL222 | ST8 | SSTI | Not known | No |
| USFL223 | ST8 | SSTI | Not known | No |
| USFL237 | ST8 | SSTI | Not known | No |
| USFL239 | ST8 | SSTI | Not known | No |
| USFL240 | ST8 | SSTI | Not known | No |
| USFL250 | ST8 | SSTI | Not known | No |
| USFL255 | ST8 | SSTI | Not known | No |

| | | | | |
|---|---|---|---|---|
| USFL256 | ST8 | SSTI | Not known | No |
| USFL258 | ST8 | SSTI | Not known | No |
| USFL273 | ST8 | SSTI | Not known | No |
| USFL274 | ST8 | SSTI | Not known | No |
| USFL276 | ST8 | SSTI | Not known | No |
| USFL277 | ST8 | SSTI | Not known | No |
| USFL279 | ST8 | SSTI | Not known | No |
| USFL282 | ST8 | SSTI | Not known | No |
| USFL319 | ST8 | SSTI | Not known | No |
| USFL320 | ST8 | SSTI | Not known | No |
| USFL326 | ST8 | SSTI | Not known | No |
| USFL327 | ST8 | SSTI | Not known | No |
| USFL330 | ST8 | SSTI | Not known | No |
| USFL339 | ST8 | SSTI | Not known | No |
| USFL341 | ST8 | SSTI | Not known | No |
